# Extracellular histones trigger disseminated intravascular coagulation by lytic cell death

**DOI:** 10.1101/2020.06.11.144683

**Authors:** Congqing Wu, Yan Zhang, Lan Li, Ankit Pandeya, Guoying Zhang, Jian Cui, Daniel Kirchhofer, Jeremy P. Wood, Susan S. Smyth, Yinan Wei, Zhenyu Li

## Abstract

Histones are cationic nuclear proteins that are essential for the structure and functions of eukaryotic chromatin. However, extracellular histones trigger inflammatory responses and contribute to death in sepsis by unknown mechanisms. We recently reported that inflammasome activation and pyroptosis trigger coagulation activation through a tissue factor (TF)-dependent mechanism. Here, we show that histones trigger coagulation activation *in vivo* as evidenced by coagulation parameters and fibrin deposition in tissues. However, histone-induced coagulopathy was neither dependent on caspase 1/11 and gasdermin D (GSDMD), nor on TLR2 and TLR4, as deficiency of these genes in mice did not protect against histone-induced coagulopathy. Incubation of histones with macrophages induced lytic cell death and phosphatidylserine (PS) exposure, which is required for TF activity, a key initiator of coagulation. Neutralization of TF diminished histone-induced coagulation. Our findings reveal lytic cell death as a novel mechanism of histone-induce coagulation activation and thrombosis.

**Key Points:**

1. Histones trigger DIC in a tissue factor dependent mechanism
2. Histones induce tissue factor activation through lytic cell death

## Introduction

Histones are essential cationic proteins that package DNA in eukaryotic cell nuclei. Extracellular histones, however, have been implicated in several pathophysiological processes.^1-3^ During innate immune response, histones released in neutrophils extracellular traps (NETs) along with DNA fibers, play an important role in entrapping and killing bacteria.^4^ Extracellular histones are highly toxic independent of NETs. Histones alone, but not NETs, activate coagulation *in vitro*.^5^ It has been reported that extracellular histones contribute to death in sepsis through TLR2 and TLR4-dependent mechanisms.^6,7^ Injection of histones into mice results in a massive prothrombotic response similar to observations in sepsis, including fibrin and platelet deposition in the lung alveoli.^7^ Coagulopathy and thrombosis are common complications in COVID-19, especially in patients who do not survive.^8-10^ Elevated histone H3 has been reported in COVID-19 and could contribute to the coagulopathy.^11^

Several mechanisms have been reported in histone-induced thrombosis. Histones can stimulate platelet aggregation *in vitro* and injection of histones into mice results in thrombocytopenia.^12,13^ Histones induce release of von Willebrand factor from endothelial Weibel-Palade bodies, which also contributes to thrombocytopenia.^14^ In addition, histones promote thrombin generation and coagulation activation through platelet-dependent and -independent mechanisms.^15,16^ Histones can increase TF activity and enhance thrombin generation in blood monocytes and endothelial cells.^17,18^ In this study, we show that histones induce macrophage lysis and phosphatidylserine (PS) exposure, leading to TF activation, which triggers systemic coagulation. Neutralizing TF by a monoclonal antibody protected against histone-induced coagulation. Our findings identify a novel mechanism of thrombosis through histone-induced lytic cell death that is independent of inflammatory receptors and signaling pathways.

## Results and Discussion

### Extracellular histones trigger coagulopathy independent of TLR receptors and inflammatory signaling pathways

Histones have been shown to increase TF activity in cells.^17,18^ To investigate whether injection of histones could directly trigger coagulation activation *in vivo*, mice were intravenously injected with unfractionated calf thymus histones at 50 mg/kg, a dose comparable to those seen in humans with sepsis/DIC.^21^ Patients with DIC exhibit increased prothrombin time (PT), elevated plasma thrombin anti-thrombin (TAT), and thrombocytopenia. Similarly, injection of histones into mice significantly increased PT (**Figure 1A**) and plasma TAT was elevated by more than 10-fold in mice that received histones (**Figure 1B**). Total platelet counts were decreased by more than 70% (**Figure 1C**). We also detected fibrin deposition in the livers of mice treated with histones, using immunostaining and western blotting (**Figure 1D-E**). Together, these data demonstrated that histones effectively trigger coagulation activation *in vivo*.

**Figure 1.**
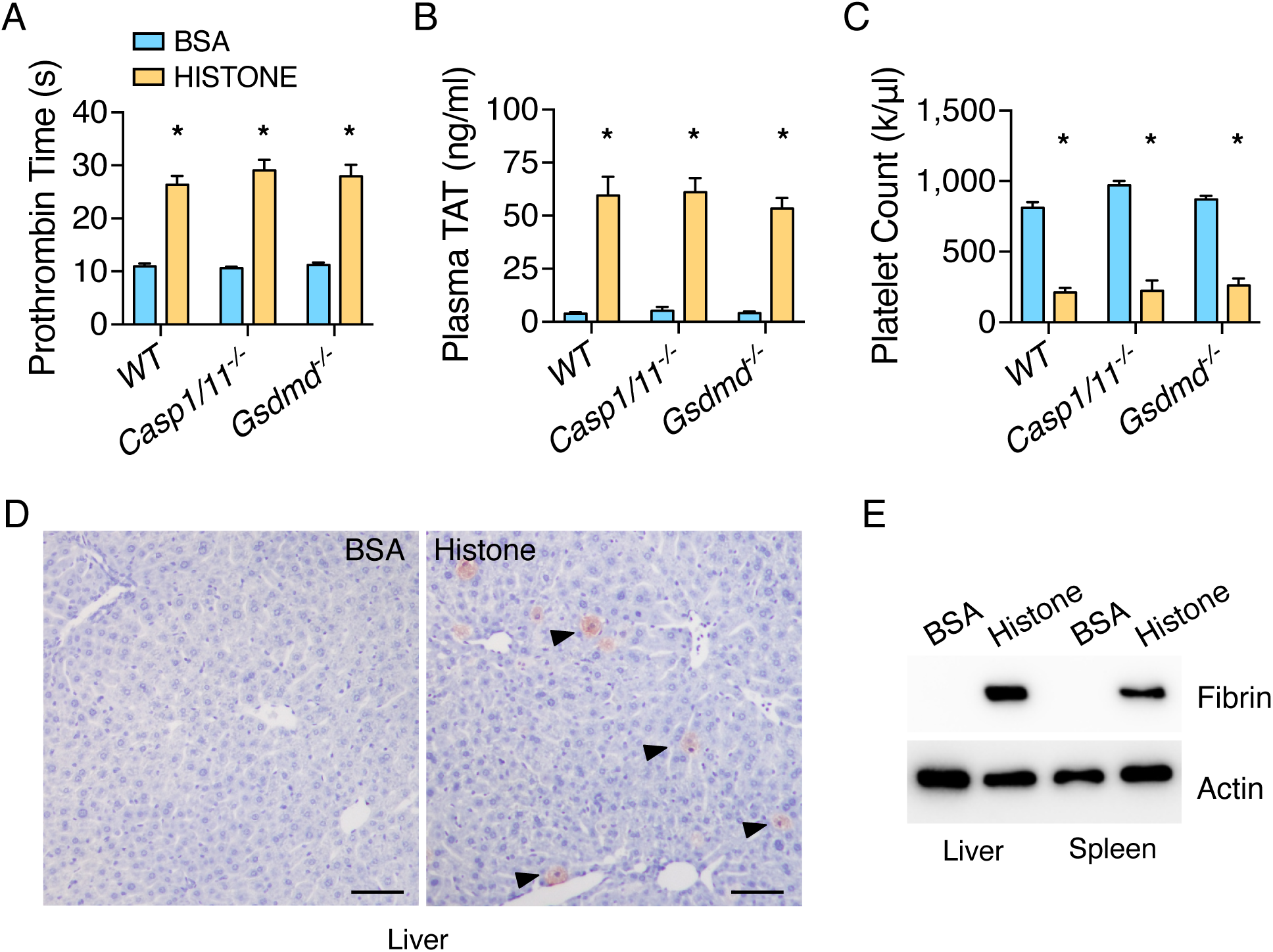
Extracellular histones trigger coagulopathy independent of inflammasome pathways. (**A-C**) C57BL/6J mice (WT), Casp1/11-deficient mice, and GSDMD-deficient mice were injected intravenously with BSA or Histones. Blood was collected 60 minutes after injection. Prothrombin time (**A**), plasma TAT concentrations (**B**), and total platelet count (**C**) were measured. Error bars denote SEM; n = 5 for all experimental groups. * P < 0.05 versus BSA, Student’s *t*-test. (**D**) C57BL/6J mice (WT) were injected intravenously with BSA or Histones. After 60 minutes, mice were perfused with PBS then perfusion-fixed with 10% formalin under physiological pressure for 45 minutes. Liver sections were immunostained with anti-fibrin monoclonal antibody (59D8). Wild type mice showed fibrin deposition in liver (arrows). Scale bars denote 50 mm. Images are representative of 3 independent experiments. (**E**) C57BL/6J mice (WT) were injected intravenously with BSA or Histones. After 60 minutes, mice were euthanized, and tissues were harvested. Fibrin in the tissue lysates was detected by immunoblot with the anti-fibrin monoclonal antibody 59D8. Data are reprentative of 3 independent experiments.

Our recent work demonstrated that inflammasome activation triggers DIC via pyroptosis.^19^ Because histones have been shown to activate inflammasome *in vitro*,^22^ we examined whether histone-induced coagulation activation proceeds through inflammasome activation and pyroptosis. To our surprise, neither deficiency of caspase-1 nor GSDMD had any significant impact on histone-induced coagulopathy measured by PT, plasma TAT, and total platelet counts (**Figure 1A-C**). This suggests that histone-induced coagulopathy is independent of inflammasome activation. Since histone-induced organ damage and death in mice involve both TLR2 and TLR4 pathways,^6,7^ we determined whether deficiency of TLR2 or TLR4 protects against histone-induced coagulopathy. Increases in PT and plasma TAT concentrations, and thrombocytopenia by histone injection were similar in wild type and TLR2- or TLR4-deficient mice (**Figure 2A-C**). These data demonstrate that histones induce coagulopathy does not require TLR2 and TLR4.

**Figure 2.**
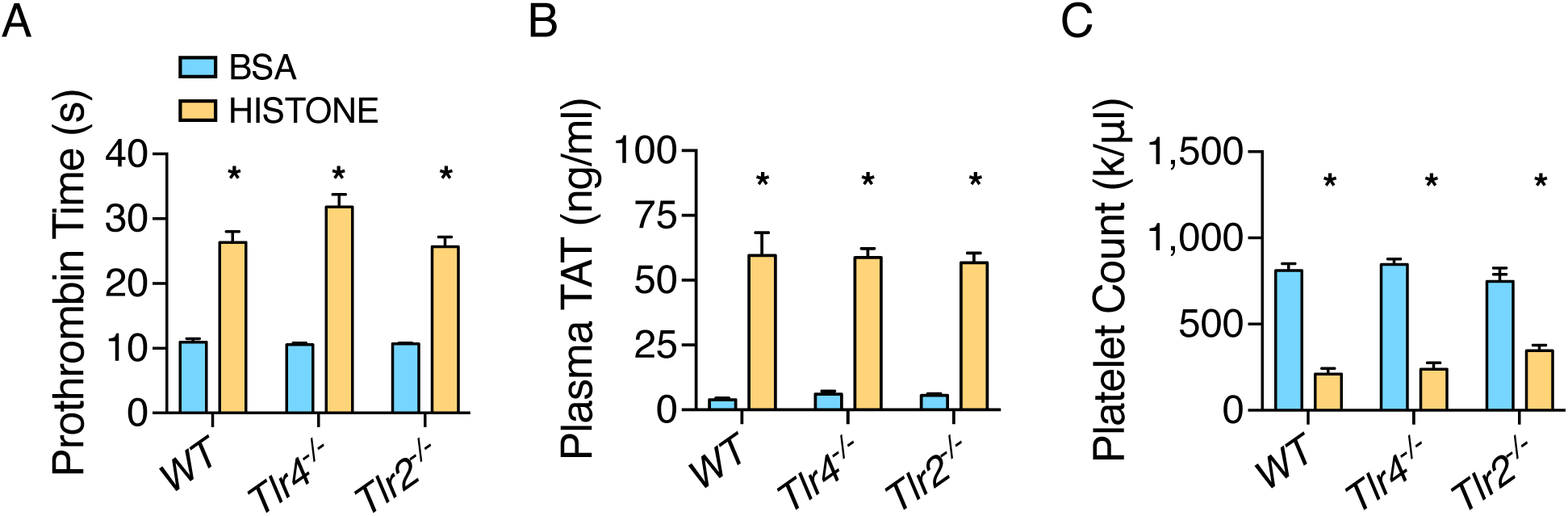
Extracellular histones trigger coagulopathy independent of TLR receptors. (**A-C**) C57BL/6J mice, TLR4-deficient mice, and TLR2-deficient mice were injected intravenously with BSA or Histones. Blood was collected 60 minutes after injection. Prothrombin time (**A**), plasma TAT concentrations (**B**), and platelet counts (**C**) were measured. Error bars denote SEM; n = 5 for all experimental groups. * P < 0.05 versus BSA, Student’s *t*-test.

### Extracellular histones lyse macrophages and expose PS

We have recently shown that TF from pyroptotic macrophages plays an important role in sepsis-associated DIC.^19^ Thus, we investigated whether histones could also induced TF activation from macrophages. Incubation of mouse bone marrow-derived macrophages (BMDMs) resulted in cell death within one hour (**Figure 3A**). Surprisingly, deficiency of caspase-1/11 or GSDMD failed to protect against histone-induced cell death (**Figure 3A**). These results indicate that histone-induce cell death proceeds through an inflammasome activation/pyroptosis-independent mechanism. In further agreement with our *in vivo* observations (**Figure 1A-C**), neither TLR2 nor TLR4 deficiency prevented histone-induced cell death (**Figure 3A**).

**Figure 3.**
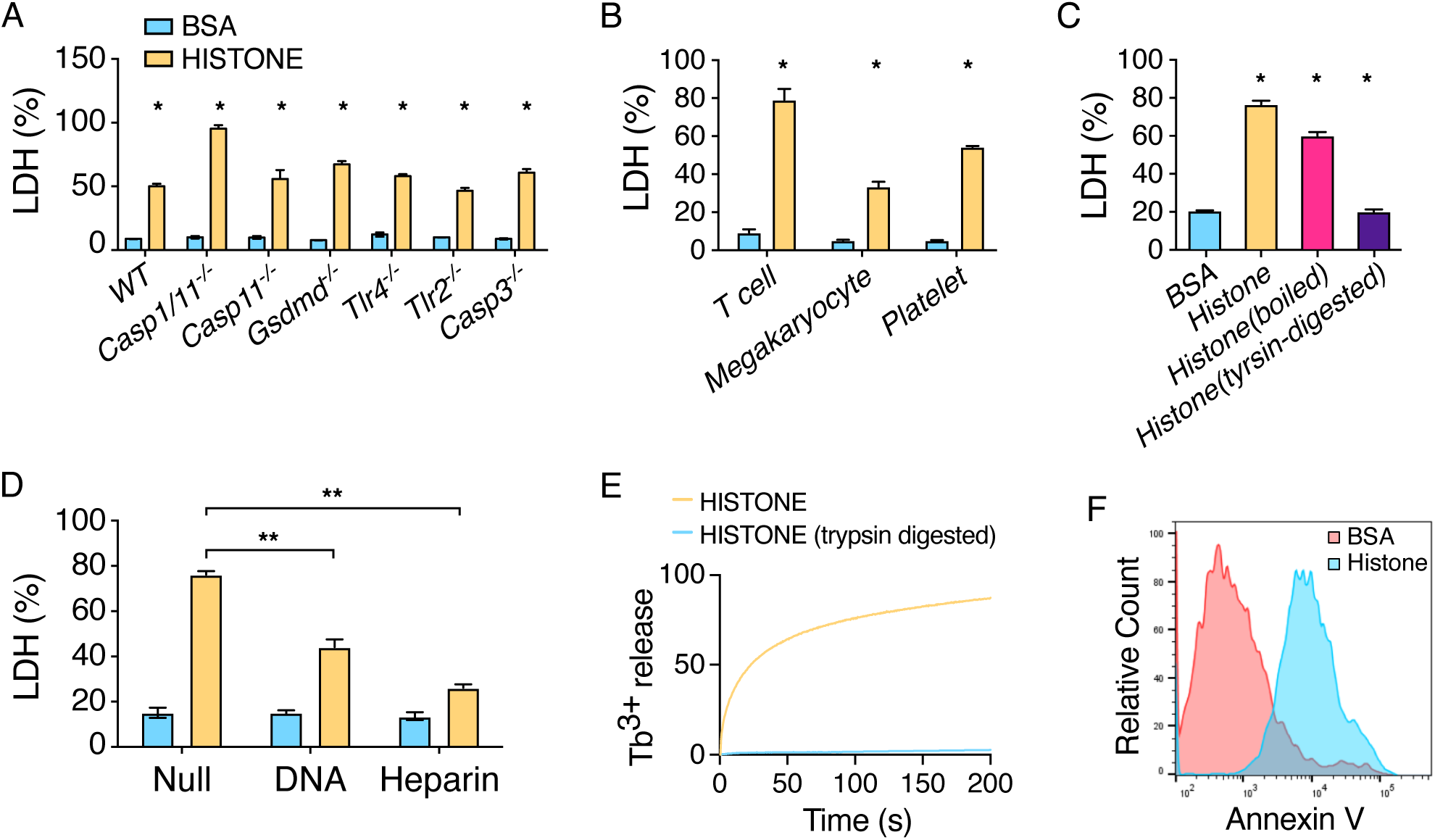
Extracelluar histones induce lytic cell death and PS exposue. (**A**) Histones lyse macrophages. BMDMs were isolated from C57BL/6J mice (WT), Casp1/11-deficient mice, Casp11-deficient mice, GSDMD-deficient mice, TLR4-deficient mice, TLR2-deficient mice and Casp3-deficient mice. Cells were incubated with BSA or Histones for 60min. (**B**) Extracellular histones lyse many other cell types. T cells, megakaryocytes, and platelets were incubated with BSA or Histone for 60 min. (**C**) Histones, even subjected to boiling, lyse macrophages. Histones, boiled histones, and trypsin-digested histones (500 ug/mL) were incubated with BMDM for 60 min. (**D**) Both DNA and heparin block histone-induced cell lysis. Mouse genomic DNA or heparin were mixed at equal amount with BSA or histone (500 ug/mL) prior to incubation with BMDM for 60 min. Histone-mediated cell cytotoxicity was determined using a LDH cytotoxicity assay. Error bars denote SEM; n = 3 for all experimental groups. * P < 0.05 versus BSA, Student’s *t*-test. ** P < 0.05, two-way ANOVA with Holm-Sidak method. (**E**) Liposome leakage was monitored by terbium (Tb3^+^) fluorescence after incubation with Histones or trypsin-digested histones. (**F**) Histone-induced PS exposure. C57BL/6J mice (WT) BMDMs were treated with BSA or Histones for 15min, and then stained with Alexa 488-labled Annexin-V and analyzed with flow cytometry.

Histones are positively charged proteins, which may allow for high affinity binding with the negatively charged cell membrane, which could lead to lytic cell death of smooth muscle cells through a non-programmed cell death mechanism.^23^ In agreement, we found that histones induced cell death in different types of cells, including platelets, megakaryocytes, and T lymphocytes (**Figure 2B**). To investigate the mechanism of cell killing, we incubated BMDMs with histones subjected to either boiling or trypsin digestion treatment. Boiled histones, but not trypsin-digested histones induced cell death in BMDMs similar to untreated histones (**Figure 2C**). To test whether histones disrupt cell integrity via its positive charges, histones were mixed with negatively charged DNA or heparin and then co-incubated with macrophages. Preincuation of histones with DNA or heparin protected against death of macrophages (**Figure 3D**), which is consistent with previous studies showing that histones as part of the nucleosome complex are not cytotoxic ^24^ and that heparin prevents histone-induced toxicity *in vivo*.^13^ We further observed that histone disrupted membranes of liposomes pre-packaged with Tb3^+^, as indicated by the time-dependent increase in Tb3^+^ release (**Figure 3E**). Leakage of the cell membrane leads to PS exposure on the outer membrane leaflet, which has been shown to be required for TF activity, the key initiator of coagulation. In agreement, incubation of BMDMs with histones did induce PS exposure, as shown by Annexin V staining (**Figure 3F**).

### TF neutralization protects against extracellular histone-induced coagulopathy

To determine whether TF is required for histone-induced coagulopathy, we utilized an inhibitory rat anti-mouse TF antibody, 1H1, to block TF activity.^25^ Indeed, mice administered 1H1, but not a control IgG were protected from histone-induced coagulopathy (**Figure 4A-C**), suggesting that TF plays an important role in histone-induced coagulopathy.

**Figure 4.**
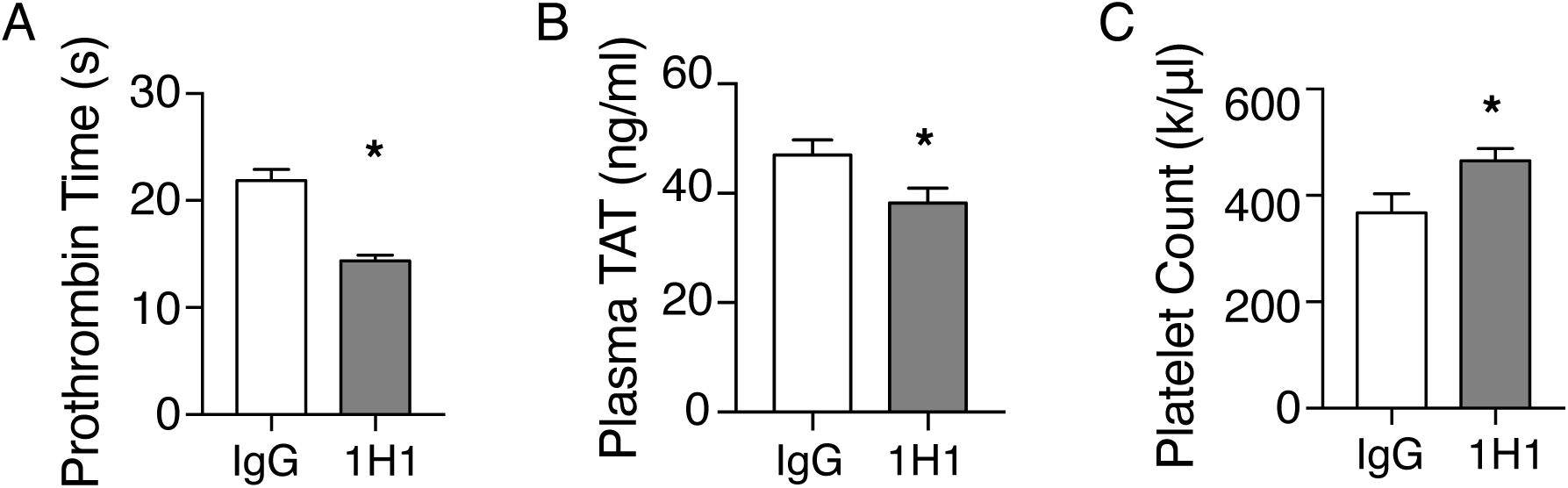
TF neutralization protects against histone-induced coagulopathy. (**A-C**) C57BL/6J mice (WT) were injected intravenously with a rat IgG or rat anti-mouse TF neutralizing antibody 1H1 (8 mg/kg). After 2 hours, mice were injected intravenously with histones. Blood was collected 60 minutes after histone injection. Prothrombin time (**A**), plasma TAT concentrations (**B**), and total platelet count (**C**) after histone injection was measured. Error bars denote SEM; n = 5 for all experimental groups. * P < 0.05 versus IgG, Student’s *t*-test.

Our results demonstrate a novel mechanism of histone-induced coagulation activation and thrombosis. In sepsis, many mechanisms contribute to DIC. However, an antibody to histones reduces the mortality of mice in lipopolysaccharide (LPS) and cecal ligation and puncture models of sepsis,^7^ suggesting that histone-induced coagulation activation is important in sepsis. Therefore, treatments that protect against histone-induced cell death may potentially increase survival of septic patients.

## Methods

### Mice

C57BL/6J, Casp1/11-/-, Casp11-/-, Tlr4-/-, Tlr2-/-, Gsdmd-/-, and Casp3-/- mice were housed in the University of Kentucky Animal Care Facility, following institutional and National Institutes of Health guidelines after approval by the Institutional Animal Care and Use Committee. Male mice at 8-12 weeks were used in all experiments.

### In vivo Challenges

Histones (Worthington cat#LS002546/lot#33P14617) were administered at 50 mg/kg via retro-obital injection.

### Pharmacological TF inhibition

Rat IgG (Sigma, Cat#I4131) or 1H1 anti-TF antibody (Genentech) at 8 mg/kg was given via retro-orbital injection 2 hours prior to histone challenge.

### Measurement of coagulation

Blood was collected from tribromoethanol-anaesthetized mice by cardiac puncture with a 23-gauge needle attached to a syringe pre-filled with 3.8% trisodium citrate as anticoagulant (final ratio at 1:10). Blood was centrifuged at 1,500 g for 15 minutes at 4° C to obtain plasma. Prothrombin Time (PT), plasma TAT Concentrations, and platelet Counts were measured as described previously.^19^ Briefly, PT was determined with Thromboplastin-D (Pacific Hemostasis, Cat#100357/lot965299) in a manual setting according to manufacturer’s instruction, using CHRONO-LOG #367 plastic cuvette. Plasma TAT levels were determined using an ELISA kit (Abcam, Cat#ab137994) at 1:50 dilution. Total platelet counts were acquired on ProCyte Dx Hematology Analyzer (IDEXX).

### Tissue Preparation and Immunohistochemistry

Mice were perfused via both right and left ventricles with PBS and then perfusion-fixed with 10% formalin under physiological pressure for 30-45 minutes. Tissues were collected and embedded in paraffin, then sectioned serially at 5 µm. Aa anti-fibrin antibody 59D8 at 4 µg/ml was used for staining fibrin deposition, with biotinylated goat anti-mouse IgG at 1:200 dilution as secondary antibody for developing positive staining.

### BMDM Cultures

BMDMs were isolated and seeded into 12-well cell culture plate at a density of 1 × 10^6^ cells/well in 1 ml of RPMI-1640 medium containing 15% L929-cell conditioned medium (LCM). BMDMs were allowed to settle overnight and refreshed with 1 ml of Opti-MEM (Thermo, Cat#31985-070) before BSA or Histones were added (500µg/ml).

### Detection of fibrin in tissues by Western blot

Frozen tissues were homogenized in 10 volumes (mg : µl) of T-PER tissue protein extraction reagent (Thermo, Cat#78510) containing cocktail inhibitor (Sigma, Cat#P8340) and PMSF. After centrifugation at 10,000 g for 10 min, supernatant was collected for beta-actin detection. Pellets were homogenized in 3 M urea and vortexed for 2 hrs at 37°C. After centrifugation at 14,000 g for 15 min, resulting pellets were suspended in SDS-PAGE sample buffer and vortexed at 65°C for 30 min. The samples were analyzed by SDS-PAGE on 4∼15% gradient gels and immunoblotted using 59D8 at 1 µg/ml.

### Liposome leakage assay

Liposome leakage assay was conducted as described^20^. Briefly, a SX20 LED stopped flow spectrometer (Applied Photophysics Ltd.) equipped with a 280 nm LED light and 400 nm cutoff filter was used to monitor the increase of fluorescence upon liposome leakage. To obtain the percentage leakage data, TX-100 0.5% (w/v) was used to dissolve the liposome and completely releases Tb3^+.^ The increase of intensity upon mixing was determined and used as the 100% leakage value F_0_. The fluorescence intensity was normalized to F_0_ to calculate percentage leakage.

### Cytotoxicity assays

BMDMs cell death was determined using LDH Cytotoxicity Detection Kit (Promega, Cat#G1780). Briefly, 50 µl of assay buffer was added to each well containing 50 µl cell culture (1 x10^5^ cells) in a 96-well plate. After incubation for 5-10 minutes, record the absorbance at 490 nm.

### Flow cytometry

Annexin V-FITC Apoptosis Detection kit (Thermo, Cat#V13241) was used to detect the PS exposure of cells in the study. Data were acquired on an CytoFlex (Beckman Coulter) and analyzed with FlowJo v10.07.

### Statistical Analysis

Data are represented as mean ± s.e.m. Student’s *t*-test (two-sided) was used to compare two-group data with normal distribution and equivalent variance; for multiple-group with two independent factors, two-way ANOVA with Holm-Sidak multiple comparisons was used for normally distributed variables. P < 0.05 was considered statistically significant.

## Authorship Contributions

C.W., Y.Z., Y.W., and Z.L. designed and performed the experiments and wrote the manuscript, assisted by L.L., A.P., G.Z., J.C., and J.S. X.L., D.K., J.W. and S.S.S provided key reagents, discussed experiments, and contributed to manuscript preparation. All authors discussed the results and commented on the manuscript.

## Acknowledgments

C.W. is a K99 awardee (K99HL145117; NHLBI). Y.W. is supported by NSF CHE-1709381, NIH, NIAID, R56 AI137020 and R21 AI142063, and NIH/NHLBI R01 HL142640 and NIH/NIGMS R01 GM132443. Z.L. is supported by NIH/NHLBI, R01HL146744 and R01 HL142640, NIH/NIGMS R01 GM132443. Dr. Wendy Katz provided help with Tissue paraffin embedding and sectioning and was supported by NIH/NIGMS Institutional Development Award P20GM103527. Dr. Hartmut Weiler at Medical College of Wisconsin and Dr. Rodney M. Camire at the University of Pennsylvania provided the anti-fibrin antibody 59D8.

